# HBP: an integrative and flexible pipeline for the interaction analysis of Hi-C dataset

**DOI:** 10.1101/083576

**Authors:** Chao He, Ping Li, Minglei Shi, Yan Zhang, Bingyu Ye, Dejian Xie, Wenlong Shen, Zhihu Zhao

**Author notes:** Email address: Chao He, Ping Li, Minglei Shi, Yan Zhang, Bingyu Ye, Dejian Xie, Wenlong Shen, Zhihu Zhao.

## Abstract

**Background:** The spatial organization of interphase chromatin in the nucleus play an important role in gene expression regulation and function. With the rapid development of revolutionized chromosome conformation capture technology and its genome-wide derivatives such as Hi-C, investigation of the genome folding becomes more efficient and convenient. How to robustly deal with these massive datasets and infer accurate 3D model and within-nucleus compartmentalization of chromosomes becomes a new challenge.

**Result:** The implemented pipeline HBP (<u>H</u>i-C <u>B</u>ED file analysis <u>P</u>ipeline) integrates existing pipelines focusing on individual steps of Hi-C data processing into an all-in-one package with adjustable parameters to infer the consensus 3D structure of genome from raw Hi-C sequencing data. What’s more, HBP could assign statistical confidence estimation for chromatin interactions, and clustering interaction loci according to enrichment tracks or topological structure automatically.

**Conclusion:** The freely available HBP is an optimized and flexible pipeline for analyzing the folding of whole chromosome and interactions between some specific sites from the Hi-C raw sequencing reads to the partially processed datasets. The other complex genetic and epigenetic datasets from public sources such as GWAS, ENCODE consortiums etc. will also easily be integrated into HBP, hence the final output results of HBP could provide a comprehensive in-depth understanding for the specific chromatin interactions, potential molecular mechanisms and biological significance. We believe that HBP is a reliable tool for the rapidly analysis of Hi-C data and will be very useful for a wide range of researchers, particularly those who lack of background in computational biology. HBP is freely accessible at https://github.com/hechao0407/HBP/blob/master/HBP_1.0.tar.gz.

## Background

Numerous light microscopy studies have shown that chromatin is tightly packaged and organized into higher-order dynamic structures, which is essential for the genome function[1–3]. Indeed, the organization of chromatin was found to play important regulation roles in many biological processes such as transcription, DNA replication, repair and even translation [4–6]. With the occurrence and rapid development of DNA-FISH[3], other super-resolution imaging[7–9] and molecular techniques, particularly newly developed 3C (Chromosome Conformation Capture)[10] and its genome-wide derivatives such as 4C[11, 12], 5C[13], ChIA-PET[14], Hi-C[15], series of Capture-C[16] etc., the study of the chromosome folding and dynamics has made great progress in recent years and the resolution achieves an unprecedented high-level[17, 18]. In fact, the resolution of Hi-C for human genome already achieved to 1-kilobase [19], which could even be reached at the 64bp theoretically as the very recent work demonstrated [20].

Combined with the numerous newly developed computational modeling and data processing methods[21], the conventional or ‘ensemble’ C-series studies of cell populations irrefutably revealed that the interphase chromatin is hierarchically and dynamic folded to form the higher-order fractal globule structure and chromosome territories, which is coincided with the results of the DNA-FISH at the single cell level in most cases[17, 18]. It also revealed the TAD (Topological Associated Domain) is the conserved structural and functional units of the chromatin folding [22, 23]. Each TAD is composed of sub-TADs and numerous loop interactions that within the same TADs are much higher than those across TADs. The different TADs could further cluster into larger-scale compartments corresponding to transcription active or inactive chromatin[24]. Although numerous different computational modeling methods such as GenomicInteractions[25], HiTC[26], hiclib[27], ChromContact[28], HiCPlotter[29], HiC-Pro[21, 30] etc. have been developed and promote the systematic, global analysis of Hi-C results, unfortunately, we still do not have an integrated general platform to analyze interactions of global and specific sequences from mapping, filtering, interaction calling, statistical evaluation and visualization of interactions automatically. Recently, Cournac and colleagues developed some program to analyze interactions between repetitive elements[31], but the program only calculate co-localization scores to evaluate genomic imprints of the 3D folding, and thus cannot be used in a single command-line manner either. The GenomicInteractions [25] is also able to analyze the interaction network of specific sites. However, it can neither process data from raw Hi-C reads, add tracks to investigate the enrichment of histones or transcription factor binding sites, nor perform the statistical analysis. HiCPlotter focuses on the association between global Hi-C interaction heatmap and other epigenetic features such as histone modification[29]. But it cannot investigate interactions between specific types of chromatin sequences. ChromContact is a web tool for analyzing spatial contact chromosomes. It can retrieve all the information of loci of interest from the UCSC browser directly[28]. However, it is incompatible with researcher’s user-supplied data, and cannot provide information such as interaction frequency or probability etc.

To fulfill the great challenges and provide an integrative resolution for Hi-C data processing, we present HBP, an open-source, easy-to-use complete pipeline to provide the tools for the sequence mapping of raw reads, extraction and normalization of chromosome interactions, heatmap visualization of interaction frequency, network analysis of specific interactions, statistical evaluation with adjustable parameters. HBP is an intact, integrative all-in-one workflow for the whole process of Hi-C data mapping, filtering and visualizing. HBP can start either from Hi-C raw dataset or other existing interaction frequency matrix. Here, we illustrate features of HBP by applying it to some publicly available datasets, and also show some interesting findings.

## Implementation

HBP is a publicly available R package for processing Hi-C datasets. It uses Hi-C raw datasets or interaction matrix files from hiclib[27] or HiC-Pro [30], a BED format file containing specific sites information. HBP is also compatible with the tracks files of histone modification or transcription factors DNA binding motif enrichment information of ChIP-seq which can be downloaded from the UCSC [32] or user-own-supplied files. Users can examine the interaction frequency distribution, explore the network topology information such as the list of name, degree, closeness, betweenness, local cluster coefficient, eigenvector centrality of network nodes etc. Besides providing the global scatter diagram to pinpoint locations of specific sites, circos[33] pictures with different tracks, network clusters according to these tracks or topological structure of the global genome and statistical evaluation, HBP could also show the frequency, number and distribution of interactions between specific sites in the genome. These information are very important for the understanding of the chromatin interaction networks. Here we provide vignettes detailing the use of HBP in Hi-C data.

### Data import

HBP can import different categories datasets such as the raw Hi-C sequencing files or the output files from HiC-Pro [30], specific sites information of standard BED format and tracks information of WIG format etc. This flexible feature allows users to easily integrate the datasets from many different sources such as ENCODE[34], modENCODE [35], GWAS [36]consortium and UCSC [32] etc.

### Interaction frequency distribution

Following the Gu’s work [37], HBP sets up and down thresholds, and discards the interaction-pairs whose frequencies are lower than the down threshold or higher than the up threshold. Then HBP separates chromatin interaction frequencies into many bins (referred to as frequency bin), and computes density for specific sites in each frequency bins. Meanwhile, HBP will compare the results with that of random control groups which were generated according to the distribution of these imported specific sites. Based on these imported specific region, every member of random control groups was calculated and made in the similar region. This comparison results make it easier for users to ascertain whether the observed interactions were biologically significant and whether these specific sites have effect on the conformation of chromatin or not.

### Interaction network topological analysis

HBP maps sequence sites of interest to the probability matrix, and calculate the interaction frequency and pinpoint the interaction networks mediated by these sites. The igraph package was used to plot an interaction network graph and make topological cluster analysis. According to the clustering results, users can classify all sites of interest into different types or clusters, and examine their potential differential properties respectively.

### Visualization of interactions and tracks

The package can visualize the interaction of interested sequences. It is compatible with the adding of some tracks such as histone modifications or transcription factors binding sites enrichment. From the graph, users can easily find out the association rules of the chromatin interactions and other genetic and epigenetic features of the interested sequences, put forward hypotheses and further design related experiments to verify.

### Statistical significance tests

HBP can also perform cluster analysis using histone modifications or transcription factors binding tracks, and make Kruskal-Wallis rank sum or T test to evaluate the statistical difference of interaction frequency, or the interaction strength of the specific sites respectively. These tests will tell us whether there exist significant differences, suggesting different properties of these sites and the interactions.

## RESULT

### Usage examples

#### Investigating effect of different repeat elements on the chromatin conformation

The repeat sequence accounts of around 50% of human genome and is reported to be related to many different physiological process of genome [34, 38]. We and others previously reported some repeat sequences are involved in chromosome interactions [11, 31, 37]. Here we examine whether all the different kinds of repeat sequence might mediate extensive chromatin interactions. To this aim, we provide a demonstration of using HBP to perform an example analysis of Hi-C data from human IMR90 cells (GEO dataset GSM1551600) [19]. With the built-in function to import data from HiC-Pro interaction file format, HBP pre-processed the Hi-C data by HiC-Pro package, and the result was normalized by the iterative correction algorithm of HiC-Pro to remove systematic noise and bias. After downloaded from Repbase in the form of FASTA file [39], and aligned to hg19 by Blat [40], the datasets of repeat elements of L1, LTR retrotransposon, SINE, satellite, ERV2 and DNA transposons etc. were inputted into our pipeline. Then the interactions frequency distribution associated with these repeat elements was estimated. To confirm the difference, HBP also set up random control groups corresponding to the distribution of different repeat elements. Figure 2 shows the distribution of six repeat elements families. While the interaction frequency of satellite and SINE involved is much lower than random controls, that of L1 is higher than random groups. These results indicated as a family, L1 but not the satellite and SINE repeat elements, is globally mediated or maintained chromatin interactions, which is agreement with a previous model [41]. Compared with control, the frequency of other elements is fluctuated, suggesting these elements maybe need to be classified into different sub-families for further accurate analysis.

**Figure 1.**
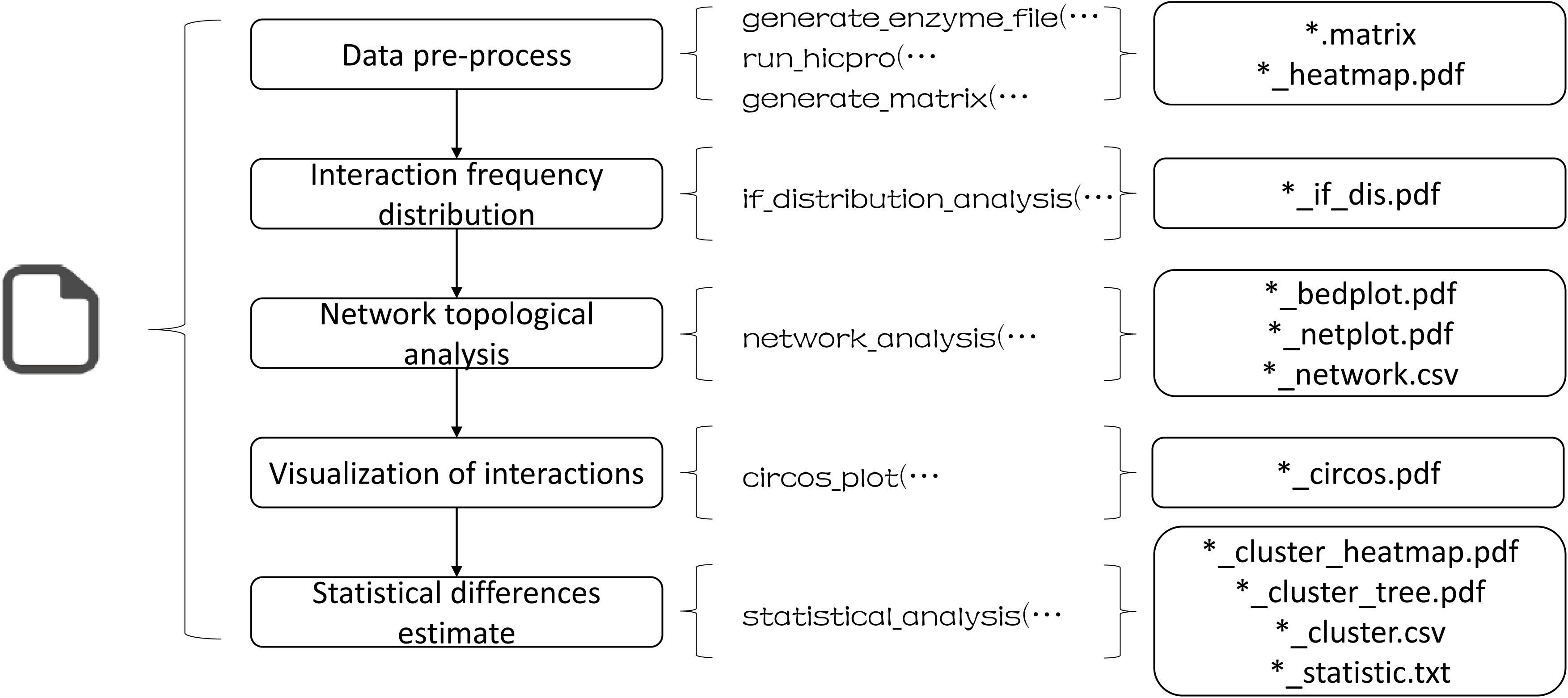
The workflow of HBP. This workflow contains the main content of the software, can be used to analyze the interaction between some specific sites.

**Figure 2.**
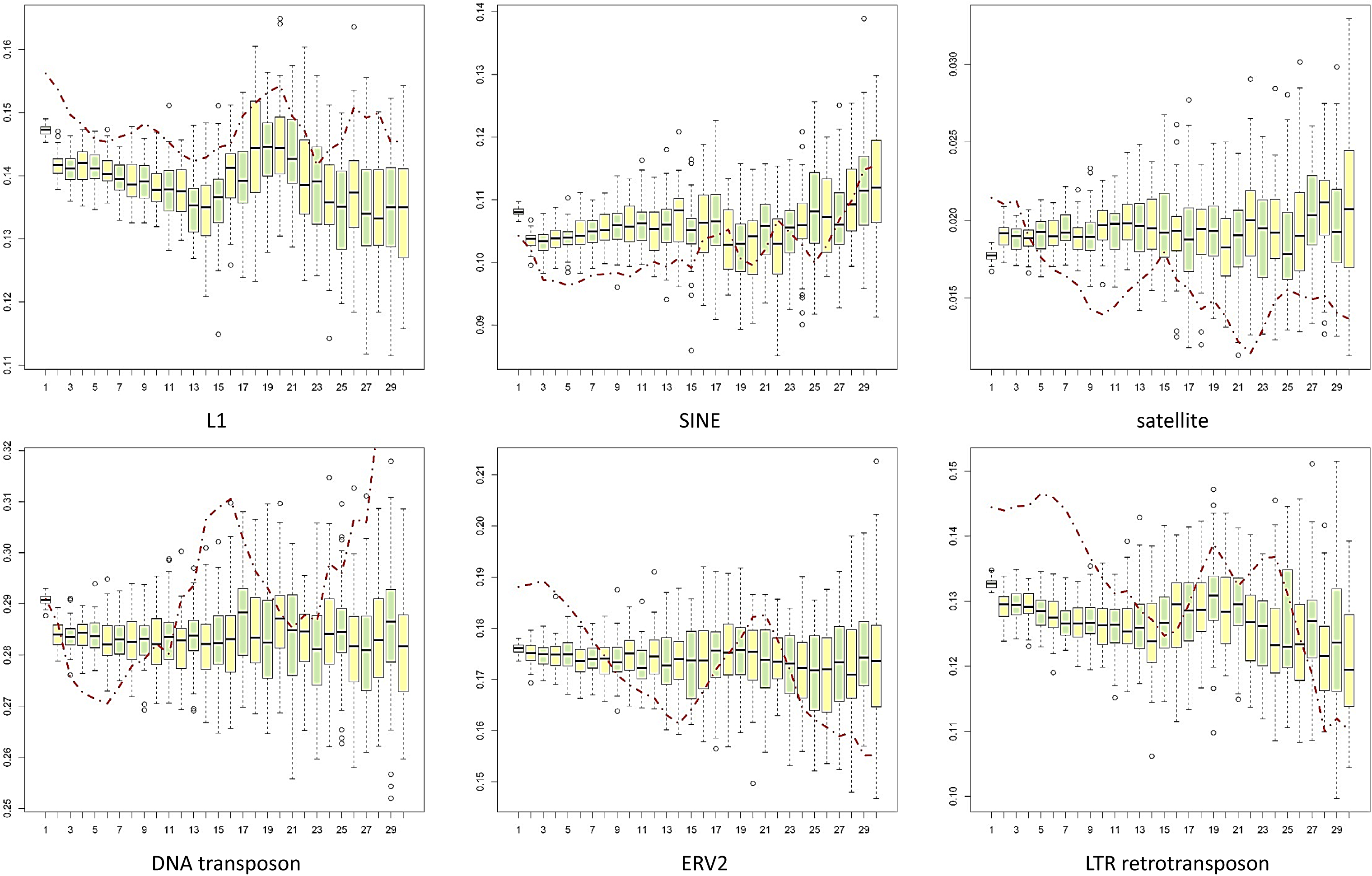
Interactions frequency distribution of different repeat elements. The red line represents interactions frequency distribution of repeat element, and the box plot represents distribution of random control.

#### Investigating the CTCF interaction network in human GM12878 cells

Alone or in combination with other protein and even RNA regulators, CTCF mediated extensive chromatin interactions [42, 43], but the topological features of CTCF-involved chromatin interaction networks remain elusive. Here we examined this issue by inputting the human CTCF DNA binding peaks of interphase chromatin into our pipeline firstly. GM12878 CTCF peaks were downloaded from ENCODE project [34, 38], GM12878 Hi-C data was downloaded from Rao’s work (GEO dataset GSM1551591) [19]. HBP processed the Hi-C dataset by HiC-Pro, invalid pairs and duplicates were removed automatically.

All CTCF-involved interactions, defined as both sites of interactions with CTCF-binding peaks around, were filtered from the dataset, and the interaction network and interaction frequency distribution were checked by HBP. As an example, we just examined the chosen region of chr8:69,300,000-74,300,000. Firstly, it was found that both the enrichments (Figure 3 a,b) and peaks (Figure 3 c,d) of the global chromosome interactions correlated with CTCF-involved specific interactions very well. These results, along with the previous comparison of ChIA-PET with Hi-C [44], strongly indicated CTCF is the main architectural protein bridging genome topology[42]. Secondly, it was found all CTCF peaks within this region (98 in total) were involved in the interaction network and have either higher or lower interactions with each other (Figure 4 a), additionally, the interaction network plotted by HBP could be sorted into two clusters (contained 46, 52 nodes respectively) according to network topological structure (Figure 4 b,c). These interaction clusters were defined as of TSC (Topological Structure Cluster), which had different distribution relative to genes position. For example, while majority of TSC1 peaks from intron, TSC2 didn’t have peaks in the 3’UTR and exons regions (Fig 4 b). The different distribution of two TSCs relative to gene position implied each of these TSCs might has different regulation function, which needs further examinations. What’s more, just like Fig 3 d showed, the two clusters just belonged to two different peaks respectively, and they were separated by the TAD boundary at chr8:71,760,000-71,800,000. Additionally, we also found the node in chr8:73,199,355-73,200,024 had the highest degree (degree = 79) and connected with more than half nodes, which suggested that this node might function as a connector or hub of the whole network. To further characterize this node, we looked into this area by UCSC genome browser. We found that it was a DNase I Hypersensitivity sites cluster with many transcription factors binding sites of EBF1, IRF4, ZNF143, RAD21, SMC3, etc. within (Figure 4 c). This result provide a potential molecular mechanism explanation for the network formation.

**Figure 3.**
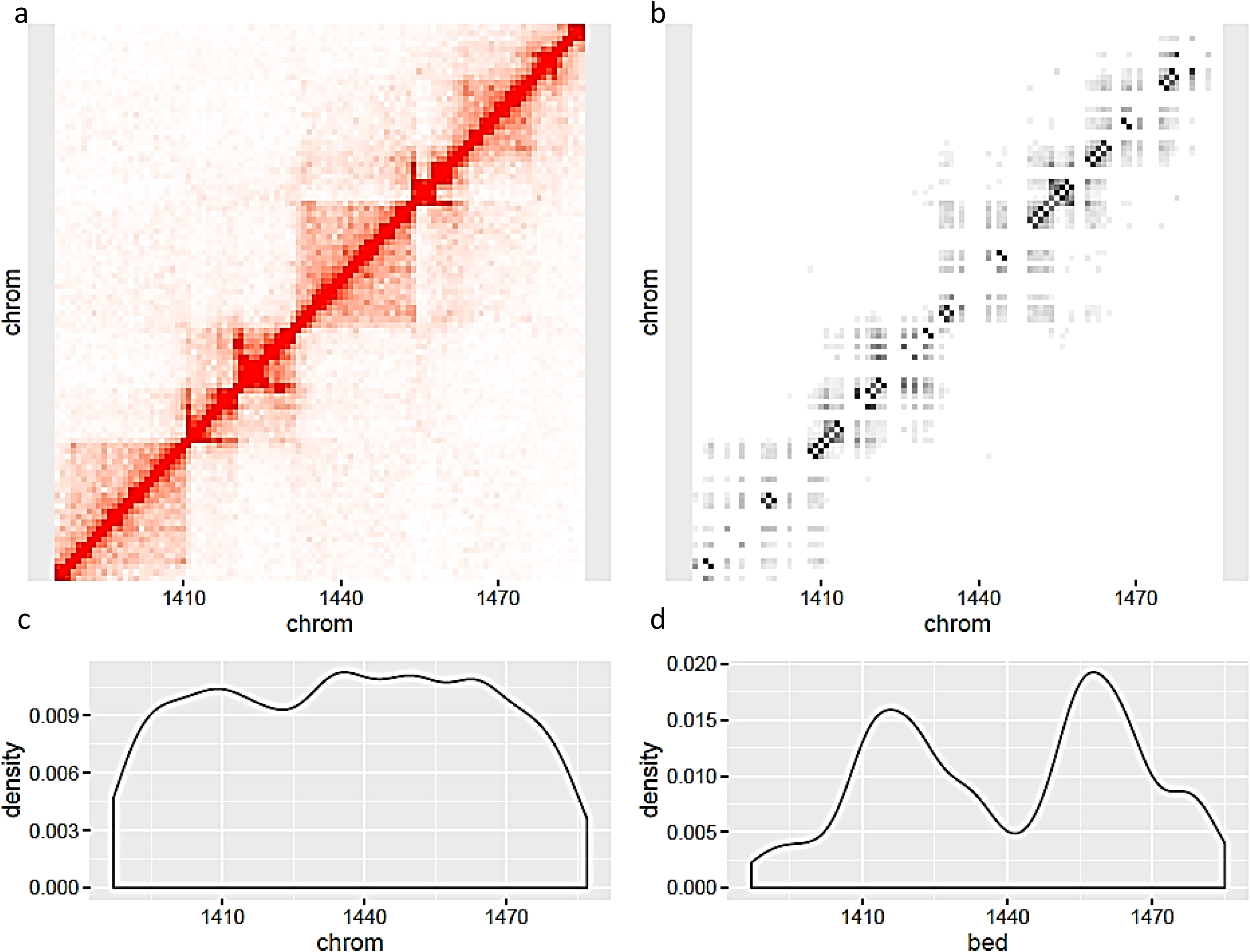
Interaction distribution of chr8: 69,300,000-74,300,000. a Hi-C interaction heatmap of this region. b CTCF involved interaction distribution of this region. c The reads counts of each bin in Hi-C interaction heatmap. d The reads counts of each bin in CTCF involved interaction distribution.

**Figure 4.**
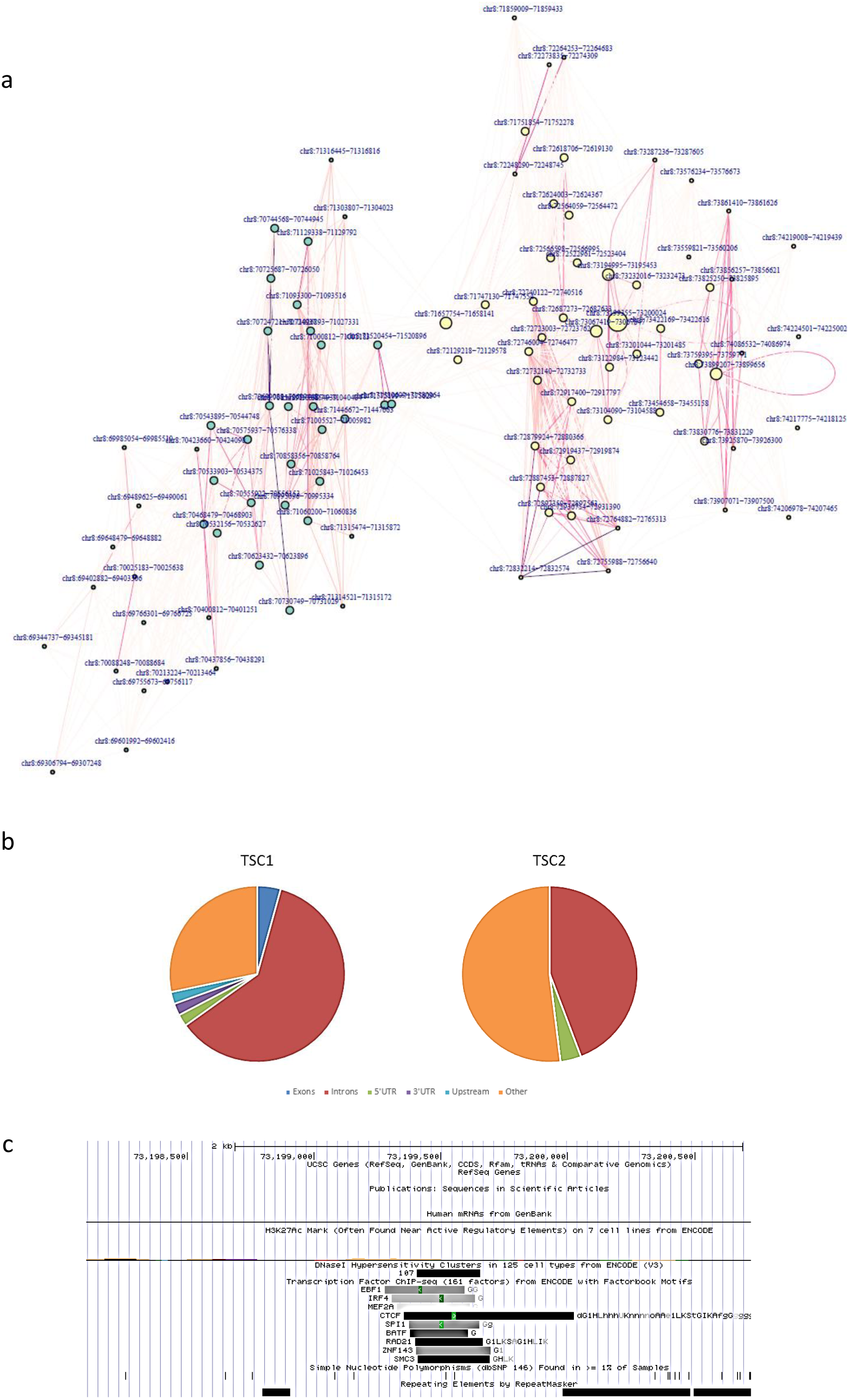
Interaction network of chr8:69,300,000-74,300,000. a The network picture without cluster. The color of interaction line represent the read counts, the size of each node represent the degree, and the color of each node represent topological structure clusters. b c The distribution of TSC1 and TSC2. The two clusters have different characteristics in relation to genes. d The highest degree site in chr8:69,300,000-74,300,000 plotted by UCSC. It is a DNase I Hypersensitivity sites cluster with many transcription factors binding sites of EBF1, IRF4, ZNF143, RAD21, SMC3, etc.

In order to learn more about the characteristics of CTCF involved interactions, we used HBP to plot a circos picture (Figure 5 a) by adding 6 histone modification features of H3K4me1, H3K4me2, H3K4me3, H3K27ac, H3K36me3 and H3K79me2. In the picture, the interaction stronger, the color deeper. We could clearly found that some interaction nodes have H3K4me1 and H3K4me2 enrichment from the tracks heatmap, which are posttranslational marks associated with enhancer regions. To know more details, all above mentioned six histone modification datasets were inputted and histone modification clusters were made by HBP, a cluster tree and a heatmap sorted by these clusters was plotted (Figure 5 b). These clusters had different modification enrichment according to the heatmap. While cluster 2 has very lower even no any enrichment of all 6 histone modification, cluster 1 and 3 have relative higher enrichment of all 6 histone modifications. Additionally, compared with cluster 3, cluster1 had higher H3K27ac enrichment but lower H3K36me3 enrichment, cluster3 was just on the contrary. As the Histone H3K27ac separates active from poised enhancers[45, 46] and higher H3K36me3 enrichment marks the active exon[46, 47], these results strongly suggested CTCF cluster 1 is active enhancer whereas cluster 3 is active expressed gene bodies, especially exons.

**Figure 5.**
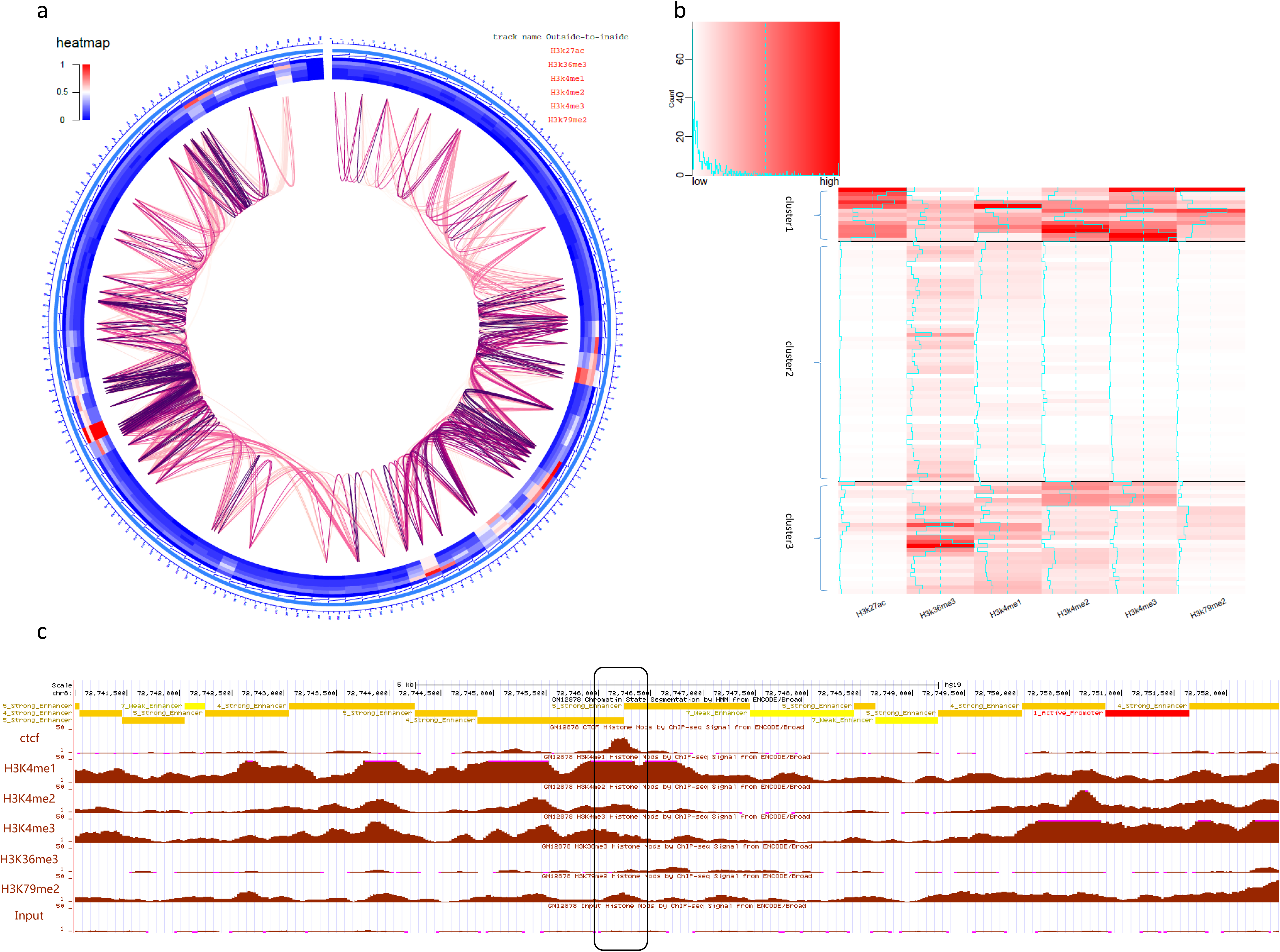
Circos plot and tracks clusters heatmap of chr8:69,300,000-74,300,000. a Circos plot of the defined region. From outside to inside, the six tracks are H3K27ac, H3k4me1, H3k4me2, H3K4me3, H3k9ac, and H3k9me3 respectively. b Tracks clusters heatmap of the defined region. The three clusters are separated by the black line in this picture. The blue line represents the value of corresponding site. c The histone modification picture plotted by UCSC. An example node from cluster 1 of b with the black square line. It is a strong active enhancer with H3K4me1, H3K4me2, H3K27ac enrichment peaks.

To further quantitatively determine whether there existed statistical differences in these histone modifications between these CTCF binding sites and other sites, Kruskal-Wallis rank sum tests were made by HBP. The test shown that H3k4me1, H3k4me2, H3k4me3, H3K27ac, H3K36me3 and H3K79me2 markers existed significant differences (p = 4.17e-05, 4.93e-03, 1.67e-07, 1.21e-03, 4.55e-05, 5.30e-04 respectively). Then T Test was carried out to estimate histone modification level of each binding site or node, and a node list containing information about modification level higher or lower than average level was obtained (Supple Table 1). What’s interesting is that 61.2% of the node shad the coherent tendency of H3K4me1, H3K4me2 and H3K27ac relative high enrichment level, which is the typical markers of active enhancers [45]. We chose one node for further analysis. It indeed just located within a super-enhancer (Figure 5 c). Super-enhancers are clusters of transcriptional enhancers that usually drive expression of genes important for defining cell identity during development[48]. What’s more, the H3K36me3 and H3K79me2 modification level in 73.5% nodes showed similar tendency, it was suggested that these site might work together in the boundary of extron-intron regions [45]. At last, T Test was also used to evaluate the interaction number between CTCF binding sites. HBP selected 100 random group with the similar distribution of CTCF binding sites to set up control. It was found that while the interaction number of random group was 1393 ˜ 1409 at 95 percent confidence interval, that of the CTCF binding sites involved in this region is 1423 (p = 1.21e-05). These results showed that in chr8:69,300,000-74,300,000, CTCF involved interaction number was much higher than random group, in other words, CTCF binding sites were more likely to interact with each other in this region.

## Conclusions

We proposed a new pipeline, HBP, to analyze global and defined interactions between specific sites from Hi-C data. Compared with other software, HBP is more optimized and flexible by providing many user-adjustable parameters, and can process Hi-C data from raw sequence or already partially processed data in an all-in-one manner. All these unique features make HBP will be a robust and useful tools for the investigation of genome hierarchical folding and functions, particularly for researchers without strong background in computational biology.

## Availability and requirements

HBP is a publicly available R package available from https://github.com/hechao0407/HBP/blob/master/HBP_1.0.tar.gz. This package requires R > = 3.3.0 and depend on several R packages including optparse, grid, lattice, IDPmisc, OmicCircos, stringr, ggplot2, igraph, reshape2, pgirmess, coin, multcomp, flexclust, HiTC, rtracklayer and gplots. All of the analyses and figures presented in the paper can be reproduced by using HBP.

## Ethics statement

Ethics statement is not applicable to our study as this study only uses publicly available data.

## Authors’ contributions

CH, WLS and ZhZ conceived the research. CH designed the software. CH, PL, MIS, YZ, WIS and ZhZ contributed to the development of software and documentation. CH is responsible for the maintenance of software. All authors contributed to the writing of the manuscript. All authors read and approved the final manuscript.

## Acknowledgements

The authors would like to thank Zhihu Zhao, Wenlong Shen, and Jing Zeng for their helpful discussions during the development of this package. This work was supported by National Basic Research Program of China [2013CB966802], National Natural Science Foundation of China [31370762, 31030026, 31272416, 81372218] and National Key Technology R&D Program of China [2012BAI01B07].

---

Supple Table 1. The list of CTCF binding sites in chr8:69,300,000-74,300,000. It contains information about degree, histone modification level, cluster membership, etc.

